# FCRLs and atypical transcriptional pattern in tumor infiltrating B cells from lung and renal cancer

**DOI:** 10.1101/2024.11.29.626090

**Authors:** EA Bryushkova, EV Shchoka, DK Lukyanov, MB Gikalo, N Mushenkova, K.K. Laktionov, A.M. Kazakov, O.A Khalmurzaev, V.B. Matveev, DM Chudakov, EO Serebrovskaya

**Affiliations:** Institute of Translational Medicine, Pirogov Russian National Research Medical University, Moscow, Russia; Department of Molecular Biology, Lomonosov Moscow State University, Moscow, Russia; Genomics of Adaptive Immunity Department, Shemyakin and Ovchinnikov Institute of Bioorganic Chemistry, Moscow, Russia; Moscow Institute of Physics and Technology; Moscow City Oncology Hospital №62, Moscow, Russia; Unicorn Capital Partners, Moscow, Russia; Center for Molecular and Cellular Biology, Russia; Federal State Budgetary Institution “N.N. Blokhin National Medical Research Center of Oncology” of the Ministry of health of Russian Federation, Moscow, Russia; Abu Dhabi Stem Cell Center, Al Muntazah, United Arab Emirates

## Abstract

Advances in high-dimensional flow cytometry and single-cell RNA sequencing (scRNA-seq) have enhanced our understanding of the heterogeneity of tumor-infiltrating B cells (TIBs). Subpopulations of TIBs exhibit diverse, sometimes opposing roles in tumor control, influenced by surface molecule, cytokine, and transcription factor expression. IgA and IgG expression in tumors have shown predictive value in melanoma and KRAS-mutated, but not KRAS wild-type, lung adenocarcinoma (LUAD).

To investigate the functional differences between B cells producing these isotypes, we performed bulk transcriptome analysis of tumor-infiltrating surface-IgA+ (sIgA+) and sIgG+ memory B cells in LUAD. In LUAD, sIgA+ B cells overexpressed FCRL4, PDCD1, and RUNX2, suggesting an atypical chronically antigen-stimulated phenotype with features of exhaustion. Public scRNA-seq data revealed FCRL4-expressing TIBs as a distinct cluster with upregulation of genes involved in IFNγ and IFNα responses.

sIgG+ B cells from LUAD overexpressed IL5RA, indicating a role for IL-5 in class-switch recombination to IgG. TCGA LUAD cohort analysis showed that the FCRL4/CD20 expression ratio correlates with lower survival, reinforcing FCRL4 as a marker of dysfunctional TIBs. Additionally, in renal cancer, high IGHA1/IGHG1 ratios were linked to worse survival.

These findings suggest that the IgA/IgG ratio in tumors reflects not only the TME cytokine environment, but also functional differences in B cell populations, providing insights into their diverse roles in tumor progression.

## Introduction

B cells may comprise more than 50% of lymphocyte infiltrate in tumors, thus being the important component of tumor microenvironment (TME)^1^. Tumor-infiltrating B cells (TIBs) exhibit the hallmarks of antigen recognition, including somatic hypermutation, class switch recombination, clonal expansion and differentiation into antibody producing plasma cells (PCs)^2^. In contrast to tumor-specific T cells that usually recognize mutated proteins (neoantigens), the target antigens of TIBs are predominantly self-proteins, thus B cells may complement the recognition pattern of T lymphocytes. All B cell subsets are found in human tumors, including activated B cells, memory and PCs, and even naive B cells.

Several reports studying the role of B cells in the context of non-small cell lung cancer (NSCLC) have shown that upregulation of B cell related genes ^3^, CD20^+^ B cell infiltration ^4^, and the presence of B cell in tertiary lymphoid structures (TLS) correlated with patient survival, suggesting B cells role in tumor control. Approximately 50% of NSCLC patients have antibody reactivity to tumor antigens ^5^. Though anti-tumor activity of B cells is well characterized, in fact TIBs may play multiple roles in pro-tumoral antigen-specific immune responses as well. Immune promoting activities of TIBs include antigen presentation to T cells; direct cellular cytotoxicity^2^, production of cytokines and antibodies. Depending on the isotype antibodies possess different effector functions. Tumor-specific antibodies may induce killing of tumor cells via ADCC ^1,6^ and complement activation, enhance antigen capture and presentation by dendritic cells^7^. Far fewer studies showed tumor promoting effects of antibodies, and mainly pro-tumoral activity of antibodies was attributed to the persistent immune complexes modulating the activity of Fc receptor-bearing myeloid cells^8^.

Antibody isotypes may be associated with distinct functional B-cell populations. IgA isotype in some models characterizes suppressive intratumoral B cell population as was clearly shown *in vivo* in the murine model of metastatic prostate cancer^9,10^. The suppressive B cell (Breg) population was described as IgA^+^CD19^+^ plasmacytes expressing IL-10 and PD-L1 as well as other immunoregulatory factors.

Bregs increased on chemotherapy and had a deleterious effect on tumoral CTL activation. The immunosuppressive IgA+TIBs are also relevant for clinical situation: based on the analysis of RNA-Seq data from TCGA, we have previously observed association of high IgA/IgG1 ratio with worse survival in melanoma patients^11^ and bladder cancer patients receiving immunotherapy ^11,12^. In case of lung adenocarcinoma (LUAD) low proportion of IgA among all intratumoral Ig was associated with improved overall survival in a specific subset of patients with mutated KRAS gene ^13^.

B cells may undergo exhaustion upon chronic antigen stimulation, a process analogous to T cell exhaustion^14–16^. This process is defined as the progressive loss of functionality and reduced proliferative capability triggered by chronic antigen stimulation. Exhausted B cells (Bexh) are most studied in the context of chronic viral infection such as HIV, and have been defined as atypical memory B cells with downregulated CD21 and CD27^14^. Phenotype of HIV-induced Bexh includes higher expressions of certain chemokine receptors (CXCR3 and CCR6) and a number of ITIM-bearing receptors that serve as negative regulators of BCR-mediated activation, including FCRL4, FcγRIIB (CD32b), CD22, CD85d, and other B-cell inhibitory receptors, such as CD72, LAIR-1 and the programmed cell death 1 (PD-1) ^14^. Among ITIM-containing receptors FCRL4 was shown to be involved in maintenance of exhausted phenotype as its knockdown results in significant upregulation of BCR-dependent proliferation and cytokine response ^16^. Exhausted TIBs with CD69^+^HLA^−^DR^+^CD27^-^CD21^-^ phenotype were identified in NSCLC patients and their presence positively correlated with the level of T regs in antigen presentation assay^17^. At present the data on their characterization in lung cancer is scarce and no publications have been found studying possible association of this functional subpopulation with prognosis of survival.

FCRL4, together with FCRL5, was first identified as a marker of a distinct B cell population in human tonsils ^18^. FCRL4+FCRL5+ cell population was shown to have undergone more cell divisions compared to other B cell subpopulations, indicating stronger or longer BCR stimulation history. Functionally, it has been shown that FCRL4 recognises J chain-linked IgA, and thus is a receptor for systemic IgA, but not mucosal secretory IgA^19^. Upon IgA binding, ITIM domain downregulates BCR signaling, reducing proliferative response by recruiting SHP-1,-2 tyrosine phosphatases ^20^. Capable of not only dampening BCR signaling, but also enhancing TLR signaling, FCRL4 acts as a molecular switch between adaptive and innate immune response ^21^. In colorectal cancer, FCRL4 together with other B cell exhaustion markers, was associated with shorter overall survival ^22^. This study also showed that FCRL4 expression correlated with such B cell markers as CD19 and CD20, indicating association with overall B cell infiltration, whereas PD-L1 was correlated with T cell markers. On the other hand, FCRL4+FCRL5+ transcriptomic signature was associated with response to immunotherapy in lung cancer ^23^.

Better understanding and characterization of exhausted TIBs populations in different tumor contexts is needed, as well as more comprehensive insights into molecular mechanisms of B cell polarization. This understanding may give us new tools for modulation of complex immune-tumor interaction and tumor control. Here we present the results of comparative bulk RNA-seq transcriptome analysis of tumor-infiltrating IgA+ and IgG+ memory B cells in LUAD tumors, and perform a comprehensive analysis of scRNA-seq of TIBs in LUAD to reveal the molecular characteristics of exhausted B cell population.

## Materials and Methods

### Human tumor specimens

A total of 8 samples of primary NSCLC were collected from NSCLC patients with stages I-III lung adenocarcinoma including both KRASmut and KRASwt tumors. 11 samples of primary renal cancer were collected from patients with all stages of clear cell renal carcinoma, undergoing surgical resection. None of the patients received any anti-cancer therapy before tumor resection. Informed consent was obtained for each participant at the N.N.Blokhin cancer research center. The study was conducted in accordance with the Helsinki declaration.

Patient demographics and tumor histology can be found in **Supplementary tables 1and 2**.

No blinding or group randomization was conducted. No prior power analysis was conducted.

### Tumor specimen processing

Tumor specimen were fragmented manually, incubated with 1 mg/ml of Liberase™ TL (Research Grade Roche, 5401020001) and 30 IU/ml DNAse I (Qiagen, 79254) in MACS Tissue storage solution (Miltenyi Biotec, 51606) and additionally homogenized with gentleMACS Dissociator (Miltenyi Biotec). Non-digestible tissue fragments were filtered with the 100-μm Cell Strainer (BD Falcon). Buffer EL (Qiagen 79217) was used for erythrocyte lysis and mononuclear cells were then separated with Ficoll-Paque™ PLUS (GE Healthcare, 17-1440-03). Single cell suspension in Versene solution with 5% human serum (Gemini Biotech) was additionally filtered with 70-μm Cell Strainer (SPL Lifesciences, 93040) to eliminate cell clumps and was subsequently used for antibody staining.

### FACS sorting

B cell populations were sorted with FACSAria III (Becton Dickinson) in RLT (Qiagen). The following antibody panel was used: CD45-Alexa700 (BioLegend Cat# 304024, RRID:AB_493761), CD3-BrilliantViolet510 (BioLegend Cat# 300448, RRID:AB_2563468), CD19-FITC (Beckman Coulter Cat# A07768, RRID:AB_2924802), CD20-VioBlue (Miltenyi Biotec Cat# 130-094-167, RRID:AB_10830096), CD38-PerCP (BioLegend Cat# 303520, RRID:AB_893313), IgG1-PE (Cytognos CYT-IGG1PE, RRID: AB_3674600), IgA-Alexa647 (SouthernBiotech Cat# 2052-31, RRID:AB_2795711).

Cells were sorted directly into the lysis buffer (RLT, Qiagen, 79216).

### Bulk RNAseq library preparation

Total RNA was extracted using TRIzol reagent (15596–018, Ambion life technologies) according to the manufacturer’s protocol, purified by AMPure-XP (Beckman Coulter, A63881). RNA pellet was dissolved in 40ul RNAse-free water (Qiagen), containing 4 IU/ul DNAseI (Qiagen, 79254) and 1 IU/ul RNAseq inhibitor (Recombinant RNAsin, Promega, N251B), and incubated for 15 min at 37°С. After DNAse treatment RNA was purified with AMPure-XP magnetic beads (Beckman Coulter) at a ratio of 1: 1. Reverse transcription and PCR amplification was performed using SMART-Seq v4 Ultra Low Input RNA Kit for Sequencing (TakaraBio, 634893).

RNAseq libraries were created using Nextera XT DNA Library Prep Kit (Illumina) with indexes Nextera XT Index Kit v2 (Illumina). The resulting libraries were sequenced with 150 bp paired-end reads on the Illumina HiSeq4000 sequencing system (Illumina Inc., San Diego).

#### Bulk RNA-seq data analysis

FastQC (RRID:SCR_014583) and MultiQC (RRID:SCR_014982) software was used to perform Quality Control (QC) of samples. The trimming of NexteraPE adapters, reads of low quality, and short reads was performed by Trimmomatic (RRID:SCR_011848).

The alignment of samples was performed using STAR (RRID:SCR_004463) with GRCh38.p13 genome assembly and GRCh38.109 gene annotation, filtering out multimappers. The quality of aligned reads was evaluated using Qualimap 2 (RRID: **SCR_026258**) software.

Gene quantification was performed using featureCounts from the Subread (RRID:SCR_009803) package. Resulting table of counts was used for differential expression (DE) analysis. Samples with zero counts, and samples with less than 50% reads aligned to exons were excluded.

Downstream analysis was done using the R programming language. Deconvolution method Xcell realized as a part of Immunedeconv (RRID:SCR_023869) package was used to exclude samples based on low level of B cell signature. For this method counts were transformed into transcripts per million (TPMs).

For DE analysis, a DESeq2 (RRID:SCR_015687) package was used. Genes for which there are less than three samples with normalized counts greater than or equal to 5 were filtered before DE analysis. Shrinkage of logarithmic fold changes (LFCs) was performed by function lfcShrink with Approximate Posterior Estimation for generalized linear model (apeglm) shrinkage estimator. To exclude correlation with isotype of B cells during further characterisation, IGH, IGK, IGL, LOC, and JCHAIN genes were removed from the analysis. Adjusted p-value less than 0.01 and absolute LFC values greater than 1 were used as thresholds to identify and visualize DE genes. For volcano plot visualization top 20 upregulated and downregulated genes were labeled using geom_label_repel extension for the ggplot2 package. For visualization of genes using a heatmap only upregulated and downregulated genes were used.

Function class scoring (FCS) method gene set enrichment analysis (GSEA) was used for the interpretation of obtained results as a part of enrichplot package. For GSEA analysis, clusterProfiler package was used with parameters minimum gene set size = 25, maximum gene set set = 500 and p-value cutoff = 0.05.

To assess the known and predicted protein interactions between target DEGs, package STRINGdb was used with parameter “medium confidence”. Upregulated and downregulated genes were considered separately, as well as together.

### Single-cell RNA seq data analysis

Processed cellranger-aligned 10X Chromium 3’ scRNAseq data of immune cells from 35 NSCLC was obtained from effiken/Leader_et_al Github repository ^24^. The data was analyzed using the R programming language, most of the analysis was done via Seurat (RRID:SCR_016341) v4. To obtain high-quality scRNA-seq data, several filtering measures were applied to the raw matrix for each cell: the thresholds of maximum number of features < 4000, minimum number of features > 200, percent of mitochondrial RNA < 20, and percent of a ribosomal RNA > 5 were used. The dataset was processed by the use of the DoubletFinder (RRID:SCR_018771) package to identify clusters with doublets. To exclude biases during clustering at all stages, MALAT1, XIST, immunoglobulin, mitochondrial genes and genes involved in cell cycle were excluded from the list of variable genes. The first clusterization was produced using 2000 variable features, 40 dimensions and the resolution of 2.5. As a result of the first stage of clusterization, 4 clusters with 16389 cells were selected for downstream analysis based on non-plasma B cell marker expression (MS4A1 and CD19), excluding clusters marked as doublets according to DoubletFinder and clusters with expression of T cell, naive B cell and plasma cell markers (CD3E, CD8A, TCL1A, IGHD, CD38). The same procedure was also reproduced for samples from normal tissues from the same dataset, resulting in 2032 memory B cells from 28 patients.

Seurat algorithm of batch correction SelectIntegrationFeatures with parameter k.weight = 15, with the exclusion of immunoglobulin, mitochondrial, ribosomal and heat shock proteins (HSP) genes was used to account for differences between methods of library preparation of the samples. Downstream clustering and dimensional DE analysis was done with a min.pct parameter = 0.05 and minimal p_val_adj = 0.001. After clustering and DE, the cluster of interest was chosen based on a differentially expressed gene of interest: FCRL4. The GSEA function from clusterProfiler package and gseaplot function from enrichplot package were used for GSEA analysis and visualisation. LFC values of DEGs of the 13th cluster were used for gene ranking. For the detailed scheme of scRNAseq data processing, see **Fig. S1A**. The quality of batch correction can be assessed on **Fig S1B**.

### TCGA data analysis

Most of the analysis was carried out using Gepia2 web server (Gene Expression Profiling Interactive Analysis 2 (RRID:SCR_026154)) ^25^ (**Fig. 4 A, B, C, E**).

For survival analysis split by stage (**Fig. 4 D**), The Cancer Genome Atlas (RRID:SCR_003193) (TCGA) bulk tumor transcriptomic data (FPKM normalized) and clinical information of 478 LUAD patients and 515 KIRC patients were downloaded from the UCSC Xena (RRID:SCR_018938) (https://xenabrowser.net/) to study FCRL4 prognostic significance. To study the association of FCRL4 expressing B cells with OS, TCGA LUAD patients were split into high and low-FCRL4 groups based on the median cut-off value. The Kaplan–Meier method was employed for survival analysis, and the log-rank test was used to determine the statistical significance of the differences.

### qPCR

For qPCR, we used amplified cDNA libraries generated by SMART-Seq v4 Ultra Low Input RNA Kit for Sequencing (see Bulk RNAseq library preparation step) with additional 10-11 cycles amplification. The obtained amplicons were quantified with Qubit fluorometer (Thermo Fisher) using dsDNA HS kit (Thermo Fisher). Concentrations were then adjusted to the same value for the qPCR step.

ABL1, GUSB, HPRT and TBP were selected as reference genes based on their levels of expression in the memory B-cells of lung adenocarcinoma patients.

Primers used for qPCR are summarized in **Table 1**.

**Table 1.**
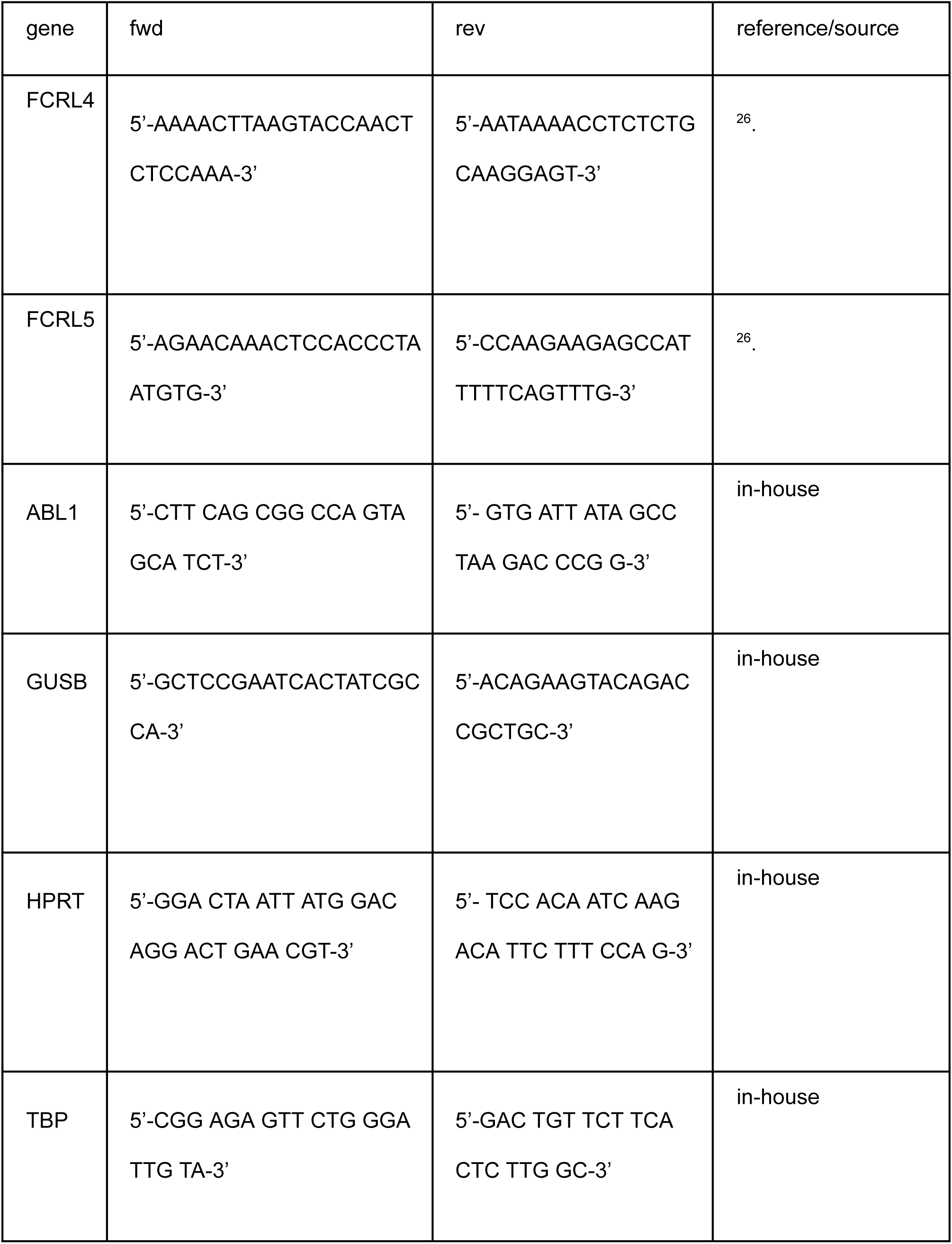
Primers for qPCR.

qPCR was carried out in 25 μL reaction volume containing 1X qPCRmix-HS SYBR (Evrogen), 0.2 μM each primer, and 2ng/ μL DNA template in a CFX96 Touch Real-Time PCR Detection System (BioRad). An initial denaturation and enzyme activation step of 2 minutes at 95°C was followed by amplification for 40 cycles at the following conditions: 15 seconds at 95°C, 30 seconds at 55.5°C, 30 seconds at 72°C. A final 5-minute extension at 72°C completed the protocol.

Melting curve analysis and the relative standard curve method performed in a CFX96 Touch Real-Time PCR Detection System (BioRad) equipped with CFX Manager™ Software (BioRad) and Precision Melt Analysis™ Software (BioRad) were used to ensure that the primers amplified the desired amplicons without forming dimers, and to calculate PCR efficiency and analytical sensitivity for each pair of primers.

Relative gene expression was calculated with the modified Pfaffl equation for multiple reference genes (https://toptipbio.com/qpcr-multiple-reference-genes/) using a custom Python script.

## Code availability

Custom scripts used for data analysis is available through the following link: https://github.com/EvgeniyShchoka/Transcriptomics-Tumor-infiltrating-MemoryBCells/tree/master

## Data availability

GEO submission https://www.ncbi.nlm.nih.gov/geo/query/acc.cgi?acc=GSE287504.

## Acknowledgements

Supported by the Ministry of Health of the Russian Federation.

## Conflict of interest

Authors declare no conflict of interest. Current affiliation of S.E.O. is Miltenyi Biotec.

## Results

### Transcriptomic profiling of IgA and IgG expressing TIBs from lung and renal cancer

Tumor infiltrating B cells (TIBs) were isolated from 8 surgically excised stage I-II NSCLC tumors. All patients were treatment-naive to exclude any influence of previous therapy. Gating strategy based on CD45/CD19/CD20/CD38/CD3 was used to define B cell subpopulations such as plasma (CD45+CD3-CD19+CD20lowCD38high) and memory B cells (CD45+CD3-CD19+ CD20highCD38low, Bmem). Bmem subpopulation was further analyzed for IgA and IgG cell surface expression. Experimental design is outlined in **Fig.1**. An example of B cell gating strategy is shown in **Fig.2A**.

**Fig. 1.**
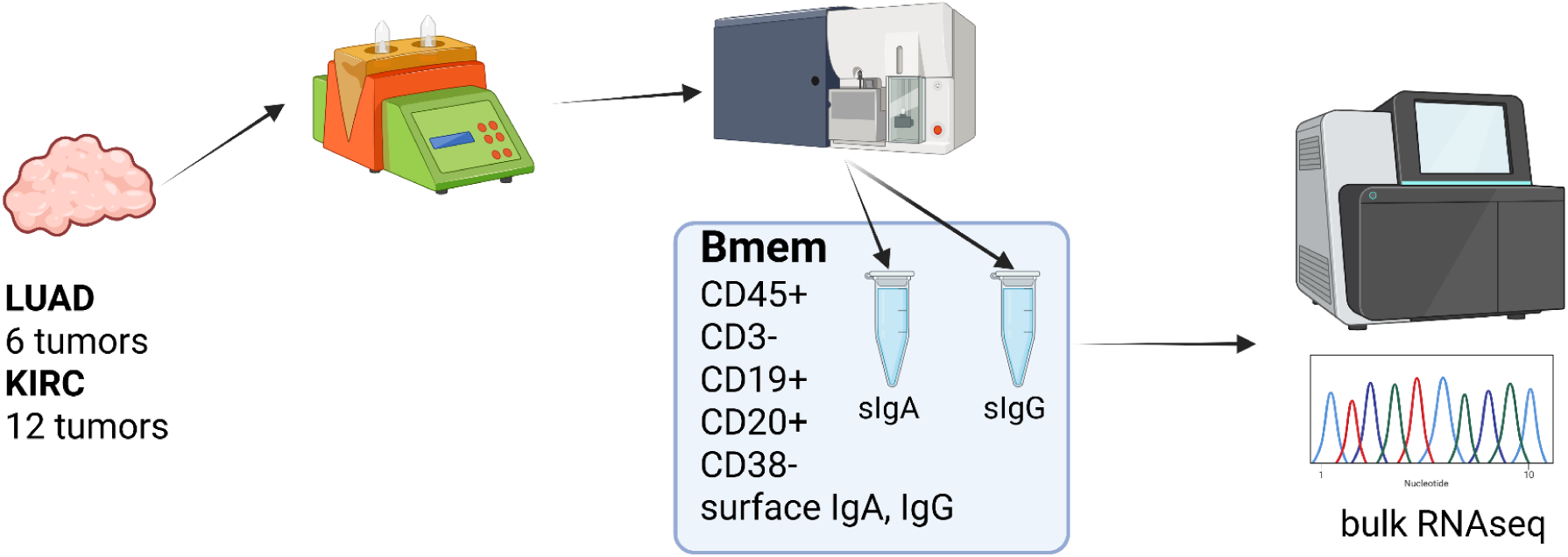
Transcriptomic profiling of surface IgA and IgG expressing TIBs, overall experimental design (Created in https://BioRender.com).

**Fig. 2.**
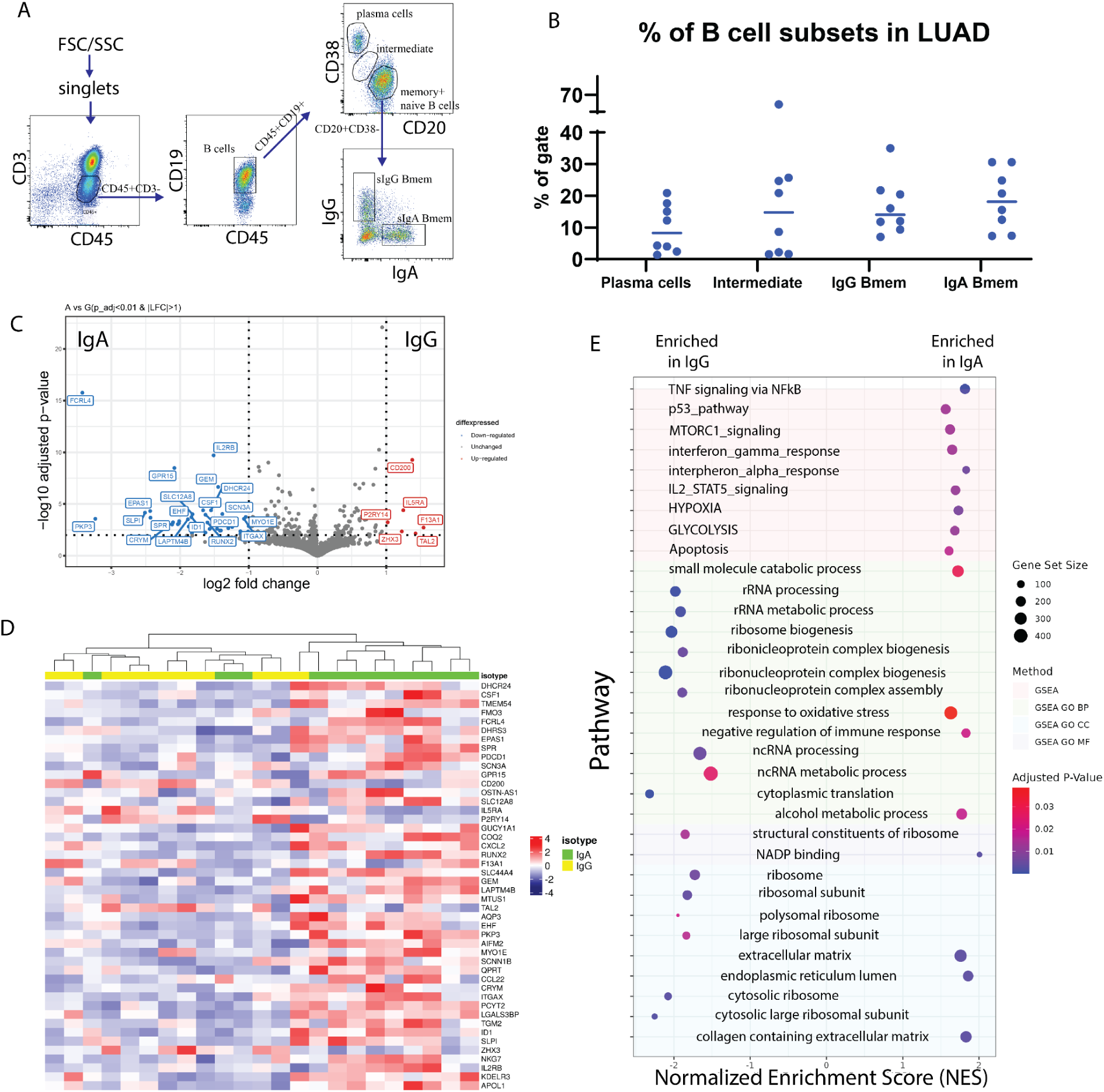
Transcriptomic profiling of surface IgA and IgG expressing TIBs LUAD. A - Flow cytometry analysis of TILs subpopulations from a representative LUAD patient (lc-p17). Subpopulations of T lymphocytes (СD45+CD3+), B lymphocytes (CD45+CD3-CD19+), plasma cells (CD20lowCD38high), (naive + memory) B cells (CD20^high^CD38^low^), IgA- and IgG-expressing memory B cells are indicated. B - Percentage of the B cell subpopulations across samples. C - Volcano plot representing differentially expressed genes in IgA Bmem (left) and IgG Bmem (right). D - Heatmap representing all differentially expressed genes. E - Gene set enrichment analysis of DE genes from panel D.

The results of subpopulation analysis (**Tables 2, 3**, **Fig.2 B**) revealed high patient-to-patient heterogeneity in the plasma to memory B cells ratio, as well as frequency of IgA and IgG-expressing Bmem cells. There was a trend for higher frequency of plasma cells in KRAS mutant tumors. IgA to IgG Bmem ratio varied from 0.3 to 15. Due to the limited number of patients it is impossible to make any conclusion about association of subpopulation frequencies and KRAS status of patients.

**Table 2.** Subpopulation composition of TIBs from LUAD patients.

| Patient | KRAS status | Plasma cells (%) | Bmemory (%) | IgA-expressing Bmemory (%) | IgG-expressing Bmemory (%) |
| --- | --- | --- | --- | --- | --- |
| lc-p7 | unknown | 22,5 | 70 | 29 | 23,5 |
| lc-p8 | wt | 1,8 | 94,2 | 19 | 15,3 |
| lc-p9 | wt | 12,8 | 83,7 | 7,3 | 35 |
| lc-p12 | wt | 5,1 | 78 | 11,8 | 10,5 |
| lc-p15 | wt | 5,4 | 67 | 7,8 | 10,7 |
| lc-p16 | wt | 5,7 | 13 | 2 | 7,2 |
| lc-p17 | G12C | 14,6 | 42 | 21 | 9,7 |
| lc-p20 | G12C | 9,3 | 63,9 | 8,9 | 18,3 |

**Table 3.** Subpopulation composition of TIBs from KIRC patients.

| **Patient** | **Plasma cells (%)** | **Bmemory (%)** | **IgA-expressing Bmemory (%)** | **IgG-expressing Bmemory (%)** |
| --- | --- | --- | --- | --- |
| rc-p12 | 13,5 | 63,3 | 71 | 5 |
| rc-p16 | 32,8 | 8 | 12 | 8 |
| rc-p17 | 0,8 | 20 | 20,7 | 11 |
| rc-p18 | 14 | 72 | 16,6 | 16,9 |
| rc-p19 | 3,4 | 75,5 | 5,0 | 1,4 |
| rc-p20 | 48 | 18 | 13 | 21 |
| rc-p21 | 4 | 10,7 | 20,3 | 12,6 |
| rc-p22 | 11 | 77 | 8,4 | 7 |
| rc-p23 | 3,6 | 89,7 | 10,7 | 36 |
| rc-p25 | 16,1 | 69,4 | 1,4 | 1,7 |
| rc-p26 | 0,65 | 95 | 0,5 | 12,7 |

Target populations corresponding to memory B phenotype defined as CD45+/CD19+/CD20+/CD38-/CD3- with either IgA or IgG expression were sorted and used for bulk RNAseq library preparation.

DESeq2 was used to identify differentially expressed genes between IgA+ and IgG+ Bmem populations. Thresholds of log2FC > 1 and corrected p-value < 0.01 were used. After shrinkage with apeglm estimator, DE analysis has shown 46 DEGs between IgA+ and IgG+ TIBs populations, out of which 5 were upregulated in IgG TIBs, and 41 were upregulated in IgA TIBs (**Fig. 2C, D**). Among the most upregulated genes in IgG high memory B cells in differential expression analysis (visualized as volcano plot, **Fig. 2C**) were **IL5Ra** (log2FC = 1.19), **CD200** (log2FC = =1.4), **F13A1** (log2FC = 1.54), **TAL2** (log2FC = 1.43), **ZHX3** (log2FC = 1.23). IL5RA is a type-I transmembrane protein, responsible for the binding of immunostimulating cytokine Interleukin 5 (IL5). Pattern of IL5RA expression is in accordance with the known functional activity of IL-5/IL-5R axis in IgH class switch and production of IgG antibodies.

Gene set enrichment analysis showed that TNF, p53, MTORC1, interferon gamma, interferon alfa, IL2-STAT5 signaling pathways are enriched in sIgA-expressing TIBs in LUAD tumors (**Fig.2 E**) as well as glycolysis, hypoxia and oxidative stress response pathways. mTOR and HIF1α belong to the central regulators of immune cells energy metabolism, important for shaping the immune response ^27^. Nutrient-limited microenvironment in the tumor niche drives B cell metabolic adaptation that may result in their functional polarization. TOR is a known inducer of HIF1α, which explains the coordinated upregulation of mTOR-HIF1α pathways. Consequently, HIF1 triggers transcriptional responses to limited oxygen and mediates the expression of genes encoding glucose transporters and glycolytic enzymes, thus shifting energy metabolism towards glycolysis as an adaptation to hypoxic conditions and increased energy demands ^27^. HIF1α is known to function in a stage-specific manner to regulate B lymphocyte differentiation and proliferation.

On the other hand, sIgG-expressing TIBs show enrichment in RNA processing, ribosome biogenesis, and other pathways related to protein synthesis.

### FCRL4 and associated transcriptional factor RUNX2 identified as overexpressed in IgA-expressing tumor-infiltrating B-cells in lung cancer

FCRL4 has been identified among genes most upregulated in IgA expressing memory TIBs. FCRL4 is a member of Fc Receptor-Like proteins (FCRL), that is able to bind soluble immunoglobulins via the Fc-domain. FCRL4 expression was previously shown to be restricted to a subpopulation of Bmem cells with a distinctive phenotype and tissue localization pattern ^28,29^. FCRL4 interacts with polymeric IgA (and to a significantly lesser extent to IgG3 and IgG4)^19^, its intracellular domain contains three consensus immunoreceptor tyrosine-based inhibition motifs (ITIMs), and co-ligation of FCRL4 with the BCR aborts BCR-mediated signaling^20^. The expression of FCRL4 was shown to correlate with transcription factor runt-related transcription factor 2 (RUNX2)^30^, which is also upregulated in IgA secreted B cells in our data (adjusted p-value = 7.36e-05, LFC =-1.65). RUNX2 gene was also shown to be specifically upregulated in CD27(-)IgA(+) B cells T-cells, independent mucosal Bmem population producing polyreactive anti-commensal antibodies^31^. In our data RNAseq, RUNX2 was upregulated in 2 out of 3 sIgA-expressing TIB samples where FCRL4 was also upregulated (**Fig. 3 A, C**). Expression of FCRL4in individual TIB samples from our cohort was further validated using qPCR and found to be significantly enriched in IgA vs IgG-expressing TIBs from LUAD (**Fig.3B**). IL5RA, on the other hand, was upregulated in the majority of sIgG-expressing TIB samples from our cohort (**Fig.3 D**)

**Figure 3.**
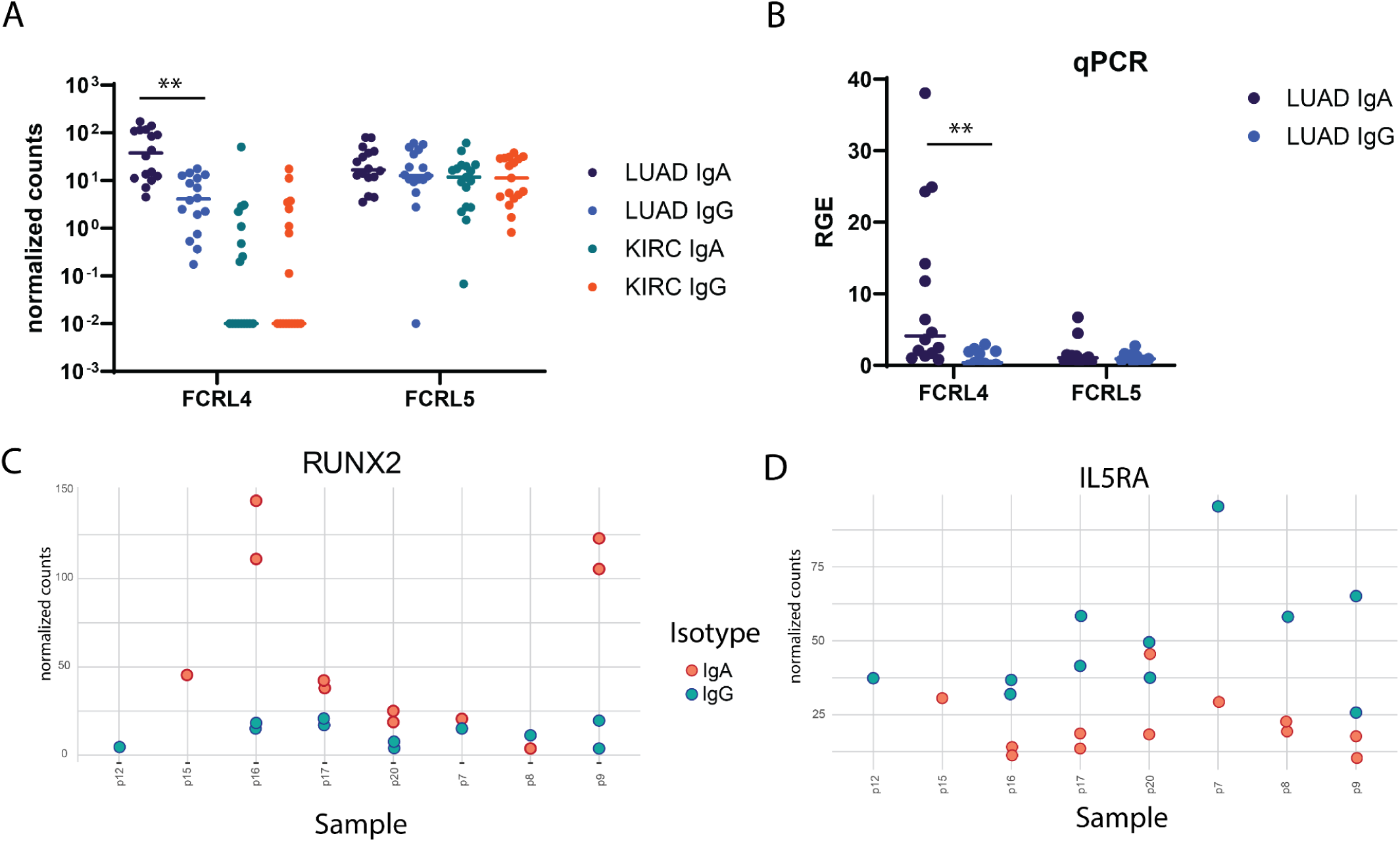
Verification of differential gene expression findings. A - per-sample distribution of expression (normalized counts) for FCRL4 gene in LUAD samples. B - Relative gene expression of FCRL4 in LUAD samples quantified by qPCR analysis. C, D - per-sample distribution of expression (normalized counts) for RUNX2 and IL5RA genes in LUAD samples (transcriptomic data).

**Figure 4.**
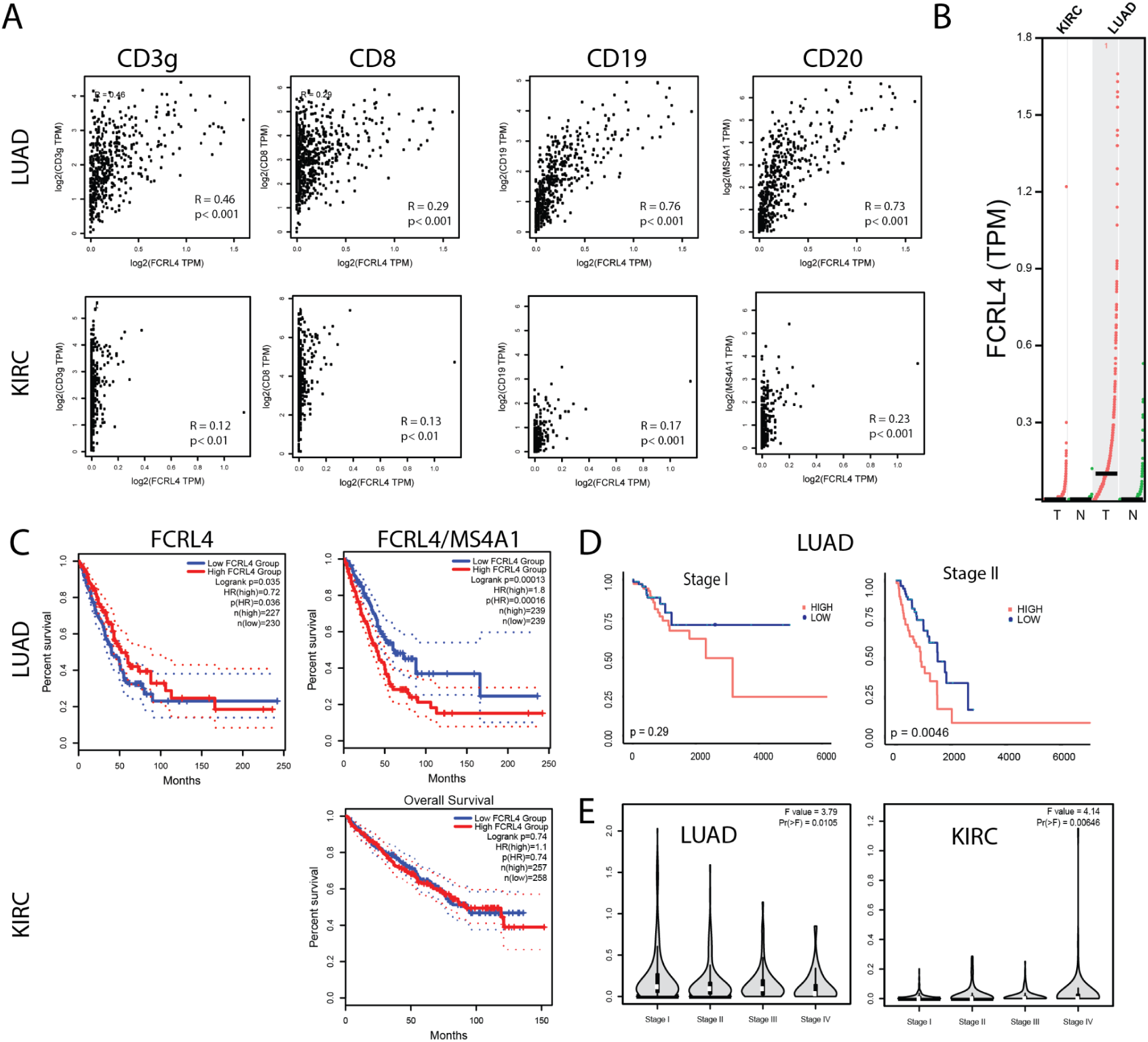
FCRL4 gene expression correlates with B cell marker expression and with patient survival. A - correlation of FCRL4 expression with CD3γ, CD8, CD19 and CD20 in LUAD and KIRC samples from TCGA database. B - FCRL4 is overexpressed in LUAD, but not in KIRC tumors versus normal tissue samples (N = 523(KIRC tum), 100 (KIRC norm), 482 (LUAD tum), 347 (LUAD norm). C - In LUAD, unnormalized FCRL4 expression positively correlates with survival (LUAD: HR_high_ = 0.72, p(HR) = 0.035), whereas FCRL4 expression normalized to B cell marker MS4A1 (CD20) (FCRL/CD20) negatively correlates with survival (LUAD: HR_high_ = 1.8, p(HR) = 0.0013, KIRC: not significant). D - For stage I LUAD, correlation of FCRL4/CD20 with patient survival is statistically insignificant (p = 0.29), whereas for stage II FCRL4/CD20 negatively correlates with survival (p = 0.0046). E - stage plots for FCRL4 expression in LUAD and KIRC. TCGA data was analyzed using the GEPIA2 server ^25^(panels A-C, E).

### FCRL4 expression normalized to total B cell infiltration is associated with decreased survival in lung cancer, but not renal cancer

The cancer genome atlas (TCGA) was used to analyze possible association between increased presence of FCRL4 among Bmem cells and survival of lung adenocarcinoma and kidney carcinoma (KIRC) patients. First, we analyzed how FCRL4 expression correlated with other immune cell markers in LUAD and KIRC samples from the TCGA database, using GEPIA2 server ^25^ (**Fig.4 A**). In KIRC, no significant correlation was observed with CD3, CD8, CD19 or CD20, and the expression of FCRL4 was overall very low. In LUAD samples, FCRL4 expression correlated well with CD20 (R=0.73) and CD19 (R=0.76). Also, in LUAD samples, expression of FCRL4 was significantly higher in tumor samples, compared to the matched normal tissue (**Fig.4 B**). In KIRC samples, FCRL4 expression was low both in tumor and matched normal tissue samples. These observations, in conjunction with our own RNAseq data, indicate that FCRL4-expressing B cells infiltrate LUAD tumors and may be functionally relevant for tumor progression. Therefore, we then performed survival analysis on the TCGA LUAD and KIRC cohorts using GEPIA2 server (**Fig.4 C**). Unnormalized FCRL4 expression weakly correlated with longer survival in LUAD (HR=0.72, p(HR)=0.035), whereas FCRL4 normalized to the expression of MS4A1 (CD20) correlated with poor survival prognosis (HR=1.8, p(HR)=0.00013). This observation suggests that whereas infiltration with B cells (reflected in total unnormalized FCRL4 expression) was a weakly positive factor, higher proportion of FCRL4-expressing B cells represented a negative factor, probably related to the higher degree of B cells exhaustion and involvement of negative feedback mechanisms of BCR signaling. In KIRC, survival data were not representative, due to low FCRL4 expression. Survival correlation with FCRL4 expression was statistically significant in Stage II LUAD patients, p=0.0046 (**Fig. 4 D**).

### FCRL4 expressing cells form a distinct cluster in tumor-infiltrating memory B cells from lung cancer, that is characterized by interferon signaling genes, antigen presentation signature and exhaustion signature

To verify the findings of our analysis of bulk RNAseq data obtained from sorted IgA and IgG TIBs from lung cancer, and also from the analysis of TCGA RNAseq data, we performed analysis of a scRNAseq dataset from Leader *et al*.^24^. As the original dataset contained CD45 positive LUAD immune cells, we filtered the data by subpopulation-specific marker expression to represent cells with memory B cell phenotype, and then performed batch effect correction and integration of data from different donors using Seurat CCA integration pipeline^32^ (**Fig.S1 A-E**).

The panel of classical B-cell markers suggests biologically relevant clusters of CD27+ memory B cells: group of IGHM+ unswitched clusters (1, 9), class-switched clusters (clusters 2, 3, 6, 8, 13), and class-switched clusters with remaining IGHM expression (0, 4, 5, 7, 11) **(Fig S1E**). Cluster 9 may belong to CR2 (CD21) high resting memory population. The marginal zone B cells (CDC1A+) did not form a cluster in this dataset.

We identified a cluster (cluster 13 on **Fig.5 A, B**) comprising 347 cells with the upregulated expression of FCRL4 (Log2FC = 1.42) relative to other clusters. FCRL4+ memory B cell cluster also has shown elevated expression of interferon response signature genes, which includes ISG15, IFIT3, IFI44L, MX2, OAS1 genes (**Fig.5 E**). Other notable genes overexpressed in this cluster include ITGAX, PDCD1, TNFSF11 and FAS (**Fig.5 C**), as well as genes associated with antigen presentation - CD83, CD86 and MHCII (**Fig.5 E**).

**Fig. 5.**
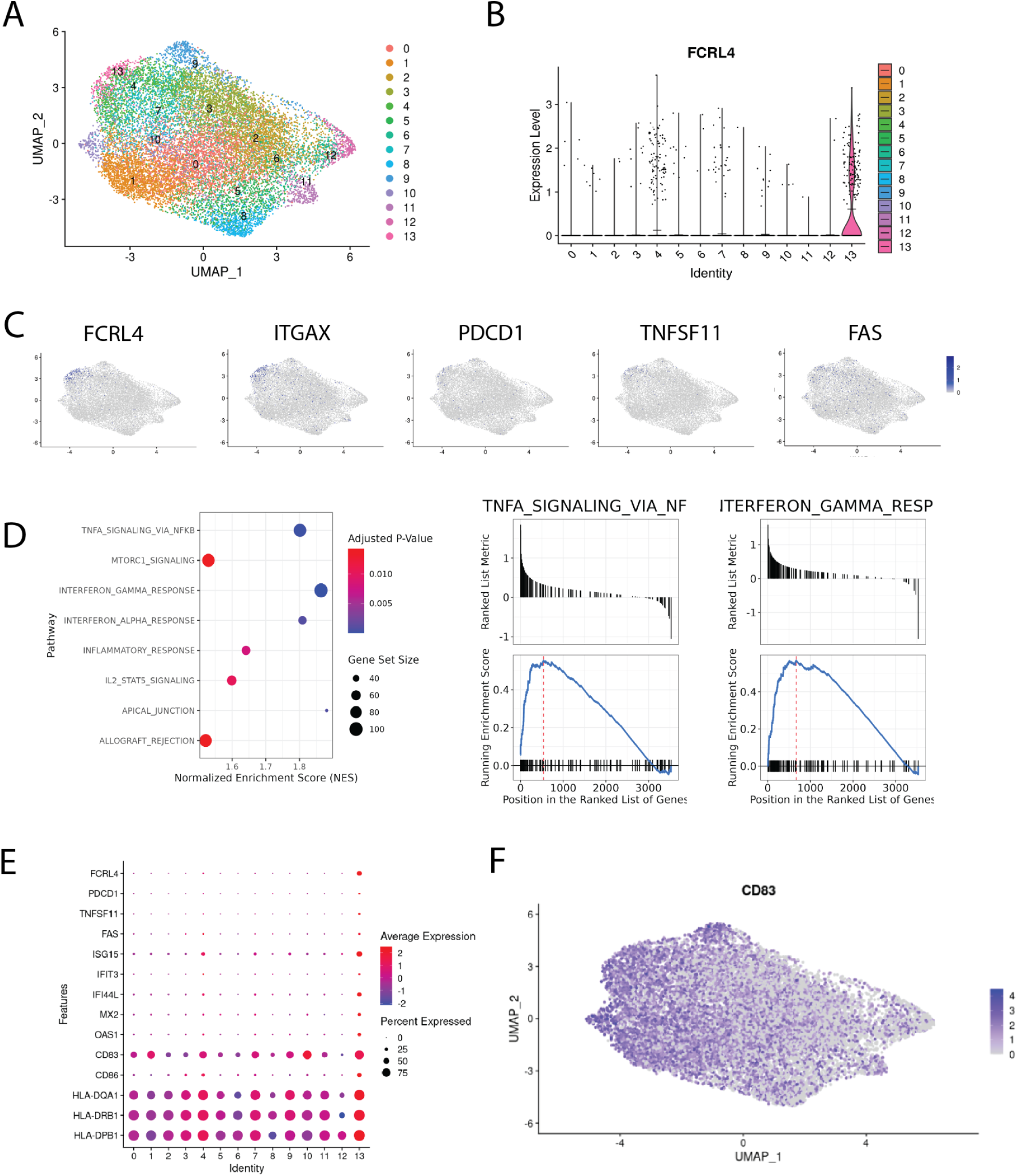
Single-cell transcriptomic analysis of memory B cells infiltrating NSCLC tumors. A - integrated Bmem dataset from ^24^. A - Uniform manifold approximation and projection (UMAP) representation of memory B cells after data integration, colors correspond to scRNAseq clusters; B - violin plot of FCRL4 expression across clusters, means are shown as black lines; C - chosen genes overexpressed in FCRL4 cluster, mapped to the UMAP representation; D - GSEA analysis of DEGs from FCRL4-overexpressing cluster 13; E - dotplot of gene expression for several cluster 13 markers across the dataset; F - UMAP representation of CD83 expression across the dataset;

ITGAX is often found in combination with other markers in the panels defining exhaustion (see ^14,33^). TNFSF11 (RANKL) was also found in FCRL4-expressing B cells in the context of autoimmunity, in the population defined as pro-inflammatory and probably contributing to RA pathogenesis^34^. Upregulation of FAS (CD95), alongside loss of CD21, was identified as a marker of B-cell dysfunction in HIV-infected individuals ^35^, contributing to B cell apoptosis ^36^.

The CD83 gene is significantly upregulated in the FCRL4-positive cluster, although not exclusively in it. The exact functions and biology of CD83 protein remain largely unknown and are actively studied as it is a clinically relevant immune checkpoint ^37^. In activated B cells and dendritic cells, membrane bound CD83 is thought to stabilize the expression of MHC II and CD86 via the negative regulation of MARCH1. In accordance with this, we find that MHC II genes and CD86 are upregulated in cluster 13 compared with other clusters (**Fig.5 E**). It was also previously shown that soluble form of CD83 promoted immunosuppression in several autoimmune conditions, restricting effector CD4 T cells and promoting Treg differentiation and proliferation. We note that both of these modes of action can potentially limit antitumor immune response, adding to the protumorigenic role of FCRL4+ tumor-infiltrating memory B cells.

We performed gene set enrichment analysis for DEGs of cluster 13, and found that IFNγ and IFNα response, IL2-STAT5, TNFa and mTORC1 signaling pathways were enriched (**Fig.5 D**). These pathways were also found to be enriched in surface IgA-expressing memory B cells from our RNA-seq data. Another set of enriched GO terms for cluster 13 DEGs is related to antigen processing and presentation via MHC class II.

Significant proportion of cluster 13 cells were found to express none or both IGHG1 and IGHA1 at comparable levels. Gating the cells according to expression of IGHA1 and IGHG1 results in roughly the same proportions of single positive IgA/IgG cells across the clusters of memory B cells (**Fig S1D**). It seems to be a general issue with isotype expression in scRNAseq memory B cell data, as in a paper by Fitzsimons et al.^38^, which included pan-cancer intratumoral B cell data from 15 different datasets, widespread coexpression of IGHA1 and IGHG1 in Bmems was also found. Thus, despite the overrepresentation of IgA-expressing cells, this limitation makes it challenging to attribute this cluster to either IgA or IgG-switched B cells. FCRL4 upregulation appears to be a tumor-specific feature, as according to the scRNA-seq data from the normal lung tissue from the same dataset, FCRL4 is not expressed by any cluster (**Fig. S2**). The study also contains CITE-seq data, however, only five B-cell relevant genes were included in the panel, IgG1 and IgA not being among them. Overall, scRNAseq data corroborates bulk RNAseq data and paints a more detailed portrait of the FCRL4+ B cell subset in LUAD.

### Low FCRL5 and high IGHA1/IGHG1 ratio correlate with prolonged survival in KIRC

To evaluate the role of B cell infiltration in renal cancer, we performed survival analysis for B cell markers on TCGA RNAseq data from KIRC samples using GEPIA2 server (**Fig.6 A-D**). All markers of B cell infiltration, including CD19, IGHA1, IGHG1, normalized to CD45, correlate with shorter overall survival, which is not the case in LUAD (**Fig.6 E-H**). Most surprisingly, the high ratio of IGHA1 to IGHG1 very strongly correlated with prolonged survival (**Fig.6 I**) in renal cancer. As isotype switching to IgA isotype is influenced by TGFβ, we hypothesized that high IgA/IgG ratio may be an indicator of TGFβ-rich microenvironment. However, neither TGFβ alone, nor TGFβ/MS4A1, nor TGFβ/CD45 ratio did not show similar correlation with survival, as IgA/IgG ratio (data not shown). Also, as opposed to LUAD, high FCRL5 expression in KIRC correlated strongly with worse survival (**Fig.6 J**).

**Fig. 6.**
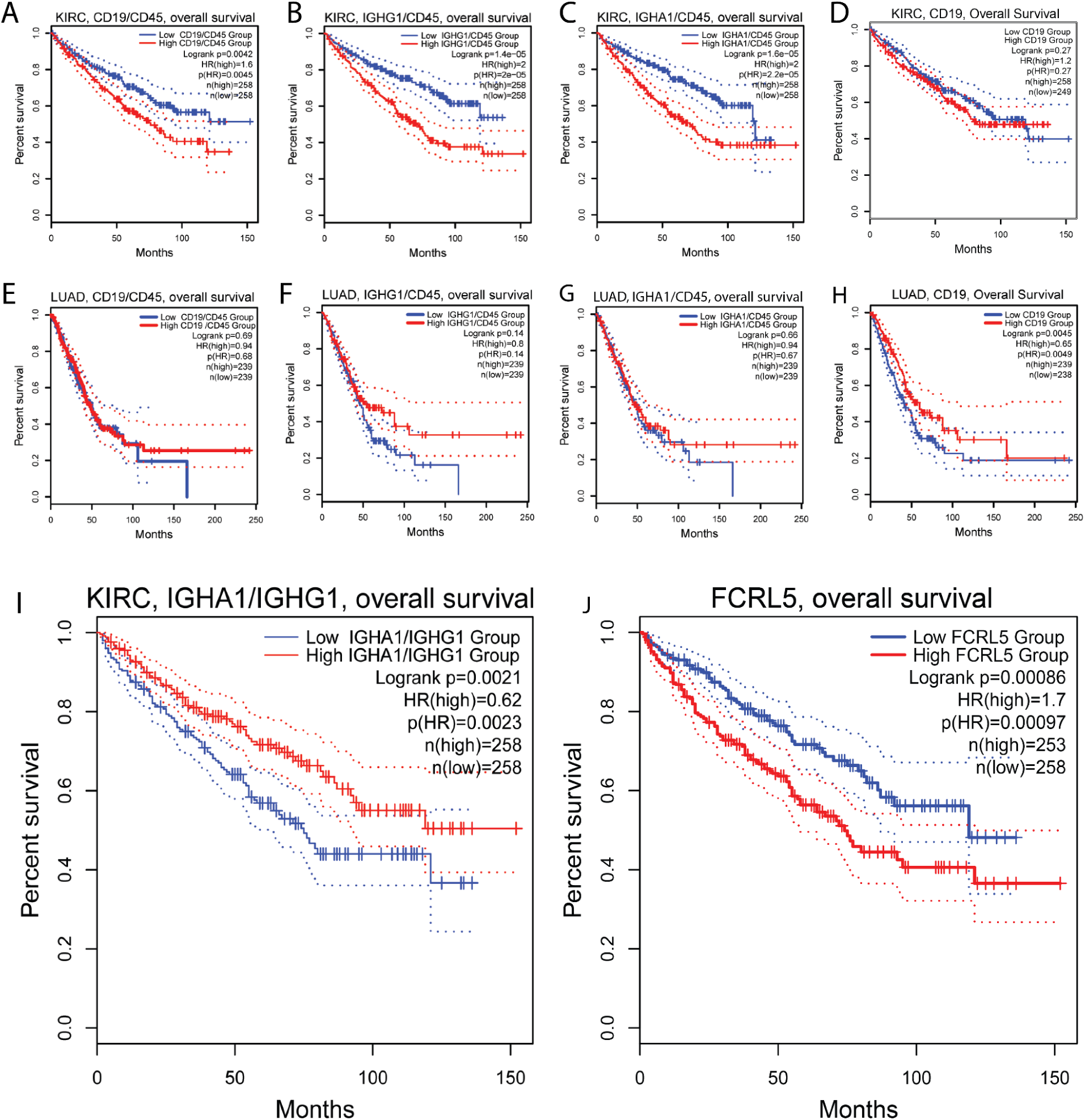
Association of B cell markers and immunoglobulin expression with survival in renal clear cell carcinoma. A-D Ratio of CD19/CD45 (A), IGHG1/CD45 (B), IGHA1/CD45 (C) are negatively associated with survival in renal cell carcinoma, indicating overall negative role of B cells in renal cancer. E-H In LUAD, no association with survival was found for ratio of CD19/CD45 (E), IGHG1/CD45 (F), IGHA1/CD45 (G); I - high ratio of IGA1 to IGHG1 is positively associated with survival in renal cell carcinoma. J - low FCRL5 expression is positively associated with survival in renal cell carcinoma.

## Discussion

The majority of recent TME research has focused on the molecular profiling of tumor-infiltrating T cells, with significantly less emphasis on the phenotypic diversity of B cells. However, it is increasingly recognized that B cells are a key component of the intratumoral immune landscape. Specific B cell subpopulations in the TME of various cancers may influence the tumor immune infiltrate, promoting either pro-tumoral or antitumoral functional profiles. B cells can amplify anti-tumor immune responses through antibody production and ADCC/ADCP. However, if they differentiate into regulatory or exhausted phenotypes, their effector function is diminished^39^. B-cell exhaustion, defined as a dysfunctional state caused by chronic antigenic overstimulation, has been observed in various lymphocyte populations^40^. It is characterized by increased expression of inhibitory receptors, abnormal chemokine and adhesion ligand/receptor expression, and weak proliferative responses to various stimuli (**Table 4**).

**Table 4.** Phenotypic characterization of Bexh population.

| Condition | Bexh phenotype | Ref |
| --- | --- | --- |
| HIV infection | CD19+CD10–CD27–CD21low | 43 |
|  | **FCRL4 high**<br>CD20highCD27–CD21low Siglec-6+,<br>CD32b high, CD22 high; CD85d high<br>CD72 high, LAIR-1 high, **PD-1 high** | 16 |
|  | CD85+ **CD11c (ITGAX) high**,<br>CXCR3 high, CCR6high , CD62 low,<br>CXCR4 low, CXCR5 low, CCR7 low | 14 |
|  | **CD95+** | 35 |
| Graft-versus-host disease | CD85+ | 15 |
| Plasmodium falciparum infection | CD85j high, CD22 high, **CD11c (ITGAX) high**, CXCR3 high; CXCR4 high, **FCRL4 high**, CD62L low, CXCR5low, CCR7low | 33 |
| Colorectal cancer | **FCRL4+**, **SIGLEC2+**, **SIGLEC6**,<br>**CD19+CD20+CD79a+** | 22 |
| NSCLC | CD69+HLA–DR+CD27–CD21– | 17 |

In this study we did not identify any chemokines or adhesion molecules differentially expressed between IgA and IgG memory B cell (Bmem) populations. However, we found upregulation of two inhibitory receptors - FCRL4 and PDCD1 (PD-1) - in IgA+ B cells in lung adenocarcinoma (LUAD). Expression of FCRL4 marks the population of exhausted tissue-like memory (TLM) B-cells previously described in HIV-infected individuals^16^. FCRL4 downregulates B-cell receptor (BCR) signaling by inhibiting BCR-mediated calcium mobilization, tyrosine phosphorylation of several intracellular proteins, and activation of protein kinase pathways^20^. Similarly, FCRL5 inhibits B cell activation via SHP-1 tyrosine phosphatase recruitment ^41^.

Considering that FCRL4 is a receptor for soluble IgA^19^, the following negative feedback mechanism may be proposed: overactivated IgA-class-switched Bmem cells sense the local IgA level via upregulated FCRL4 receptor and downregulate BCR signaling. In our current study we did not specifically measure soluble IgA production by the cells sorted as sIgA+/sIgG+ cells with our gating strategy, and it is not possible to distinguish between soluble and membrane IgA based on transcriptomic data, so currently we are unable to discriminate between autocrine and paracrine mechanisms, or a combination of both. To our knowledge, this is the first time upregulation of FCRL4 molecules in sIgA-expressing Bmem cells has been reported. We aimed to also confirm this finding with scRNA-seq analysis, but this was not possible due to technical constraints. To further explore this question, it would be beneficial to combine scRNAseq with CITE-Seq to label surface IgG/IgA, which could resolve the issue of ambiguous mapping of immunoglobulin isotypes in scRNA-seq data.

In addition to FCRL4, other markers possibly associated with B-cell exhaustion were upregulated in the IgA+ Bmem population, including PD-1, which aligns well with the B-cells exhaustion pattern described previously in chronic infections^14^.

Previously, Bexh, along with exhausted T and NK cells, were found in colorectal carcinoma (CRC) samples ^22^, where high expression of FCRL4, as well as other exhaustion genes, was significantly associated with worse prognosis independently of tumor’s molecular subtype. The overall survival (OS) of patients bearing CRC with high FCRL4 expression was more than three times shorter compared to patients with low expression. Combination of high FCRL4 and high CD20 expression was also associated with shorter OS. FCRL4+FCRL5+ Bmem population was previously identified in naïve NSCLC patients undergoing neoadjuvant immune checkpoint blockade (ICB) + chemotherapy^23^. In these FCRL4+ cells interferon-stimulated genes (CCR1, STAT1, and GBP4) and co-stimulatory molecules (CD86, СD40L) were highly expressed. Based on their gene expression profile, TLS localization and association with better prognosis and response to immunotherapy, this B cell subpopulation was proposed to possess potential anti-tumor activity that was released by ICB therapy. FCRL4+FCRL5+ B cell signature was also a good prognostic factor in melanoma^23^. Interestingly, IFNα and tumor necrosis factor (TNF) were predicted as possible drivers of FCRL4+FCRL5+ phenotype. Similarly, in our study we also observed enrichment of IFN and TNF signaling in the FCRL4-overexpressing B cell subpopulation. Immune activation proceeds in parallel with the induction of negative feedback mechanisms, which, in case of chronic inflammation, can become predominant and contribute to immune dysfunction. Type I interferons induce many of the suppressive factors that limit immunity^42^, and one effect of IFNs may be the upregulation of programmed death-ligand 1 (PD-L1), which aligns well with the observed PD-1^high^ phenotype of FCRL4+ B cells^42^.

Our current study further confirms the potential utility of FCRL4 as a prognostic marker, when used in combination with other B cell markers, and also as a potential therapeutic target. We demonstrated that FCRL4 expression, normalized to the expression of CD20, correlates with poor survival outcomes in LUAD. In KIRC, however, survival data were not conclusive, due to overall low FCRL4 expression.

Furthermore, in LUAD, we observed inverse correlation between FCRL4 expression and clinical stage. According to The Cancer Genome Atlas (TCGA) data, LUAD tumor progression was accompanied by a decrease in FCRL4 levels, whereas renal carcinoma showed a substantial increase in FCRL4 expression as the tumor progressed from stage I to stage IV. Since FCRL4 upregulation may represent a mechanism of B cells autoregulation by circulating antibodies, we suggest that increased expression of FCRL4 indicates an active humoral response. Conversely, its decrease during tumor progression may indicate the selection of low-immunogenic tumor variants that evade effective humoral control.

Notably, there is a reported trend in NSCLC patients for decreased autoantibody responses in the course of tumor progression. An analysis of the humoral response to a panel of seven autoantigens (**TP53, c-Myc, HER2, NY-ESO-1, CAGE, MUC1, and GBU4**) in NSCLC patients revealed that 71-100% had elevated antibody levels to at least one of the antigens^44^, with the highest frequencies observed at earlier stages. A similar trend was observed for anti-TOP2A and anti-ACTR3 antibodies, which showed a significant decrease in positivity from early to late tumor stages (43-50% vs. 22-24%)^45^.

Class-switch recombination to IgA isotype is induced by TGFβ^46^, an immunosuppressive cytokine produced by tumor cells, as well as by various cellular components of tumor stroma and immune infiltrate. NSCLC TME is characterized by the abundance of cancer-associated fibroblasts producing TGFB1^47^ and frequency of IgA+ cells was found to positively correlate with frequency of Tregs and exhausted CD8 T cells. Similarly, Bruno *et al.* showed that exhausted TIBs were associated with a regulatory (FoxP3+CD4+) TIL response in co-culture experiments ^17^. On the causal side, TIBs, including Bexh, may actively contribute to the conversion of T cells into a regulatory phenotype by presenting antigens to CD4+ TILs in the immunosuppressive, TGFβ-rich environment. Therefore, possible association of FCRL4+ IgA+ Bmem with Treg infiltration, as well as the degree of T cell exhaustion, presents an important avenue for future research.

Identified pattern of genes upregulated together with FCRL4 overlaps with the list of differentially expressed genes in FCRL4+ Bmem from human tonsils, with RUNX2 and IL2RB identified in both studies. Runx2 belongs to a family of transcriptional factors described in the context of early hematopoiesis and thymopoiesis ^39^. In the hematopoietic system, RUNX2 has been studied mostly in plasmacytoid dendritic cells (pDCs), in which RUNX2 facilitated pDC homeostasis and anti-viral responses. Recent data suggest participation of Runx2 in functional differentiation of T cells (Tfh, CD8 T mem) ^48–50^. Though increased RUNX2 expression is associated with various B cell-derived malignancies, including myeloma, B-cell lymphoma and acute lymphoblastic leukemia, evidence for RUNX2 role in TIBs is lacking. FCRL4 promoter contains eight potential RUNX binding sites ^30^. RUNX2 dependent regulation of FCRL4 has not been confirmed in reporter experiments, their failure may be explained by the absence of right co-factor transfection, thus the question of RUNX2-dependent regulation of FCRL4 needs adequate investigation. IL2RB represents the beta subunit of IL-2 receptor that is present in high and intermediate affinity receptor types. IL2 is a well known T cell pro-proliferative cytokine. In human B lymphocytes it was shown to imprint naive B cell fate towards plasma cell differentiation ^33^.

In this work we gained additional evidence in line with the previous reports that IgA-and IgG-class-switched B cells differ in their functional polarization and may have distinct roles in TME, important for tumor progression and survival prognosis. Our work also further supports the view that an autocrine negative feedback loop on immunoglobulin production develops throughout the course of tumor development, downregulating the activity of the B cells involved in the antitumor immune response through the interaction of inhibitory FCRLs with immunoglobulins these B cells produce.

These findings may open new avenues for research in TIBs biology, and for development of innovative immunotherapeutic strategies.

## Supplementary materials

**Supplementary table 1.** Patient characteristics - lung cancer.

| **№** | **patient ID** | **histology** | **stage** |
| --- | --- | --- | --- |
| 1 | lc-p7 | adenocarcinoma | T3N0M0 |
| 2 | lc-p8 | squamous cell carcinoma | T3N0M0 |
| 3 | lc-p9 | adenocarcinoma | pT2N3M0 |
| 4 | lc-p12 | adenocarcinoma | T2aN0M0 |
| 5 | lc-p15 | adenocarcinoma | T2aN0M0 |
| 6 | lc-p16 | adenocarcinoma | T2aN0M0 |
| 7 | lc-p17 | adenocarcinoma | T1N1M0 |
| 8 | lc-p20 | adenocarcinoma | T1BN0M0 |

**Supplementary table 2.** Patient characteristics - renal cancer.

| **№** | **patient**<br>**ID** | **histology** | **stage** |
| --- | --- | --- | --- |
| 1 | rc-p12 | renal cell clear cell carcinoma | cT3cNxM1 |
| 2 | rc-p16 | renal cell clear cell carcinoma | cT3aNxM0 |
| 3 | rc-p17 | renal cell clear cell carcinoma | no information |
| 4 | rc-p18 | renal cell clear cell carcinoma | cT3bN0M0 |
| 5 | rc-p19 | renal cell clear cell carcinoma | cT3cNxMx |
| 6 | rc-p20 | renal cell clear cell carcinoma | cT2N1Mo |
| 7 | rc-p21 | renal cell clear cell carcinoma | no information |
| 8 | rc-p22 | renal cell clear cell carcinoma | pT3aN0M0 |
| 9 | rc-p23 | renal cell clear cell carcinoma | T3cNxM0 |
| 10 | rc-p25 | renal cell clear cell carcinoma | cT2N0M1 |
| 11 | rc-p26 | renal cell clear cell carcinoma | cT3N0M0 |

**Supplementary Figure 1.**
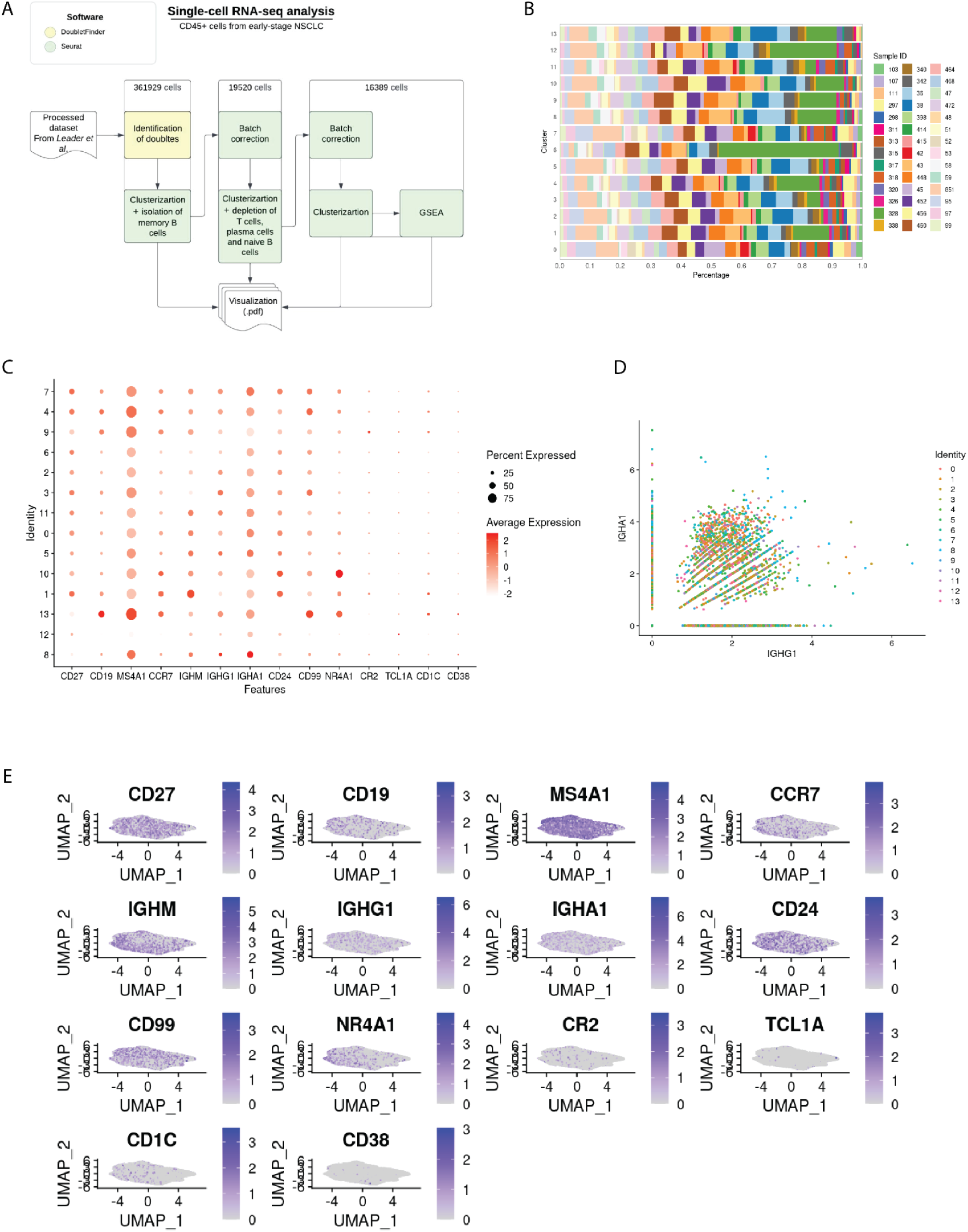
A - Diagram representing the scRNAseq data analysis steps. B - Histogram of clusters split by sample of origin to assess the efficiency of batch effect removal. C - Dotplot of B-cell subpopulation marker gene expression across 13 Bmem clusters. D - Histogram of the IGHA1/IGHG1 isotype expression across the clusters. The cells with the zero expression of one of IGHA1/IGHG1 were considered single-positive. E - B cell subpopulation marker gene expression mapped to UMAP representation of Bmem from LUAD tumors.

**Supplementary Figure 2.**
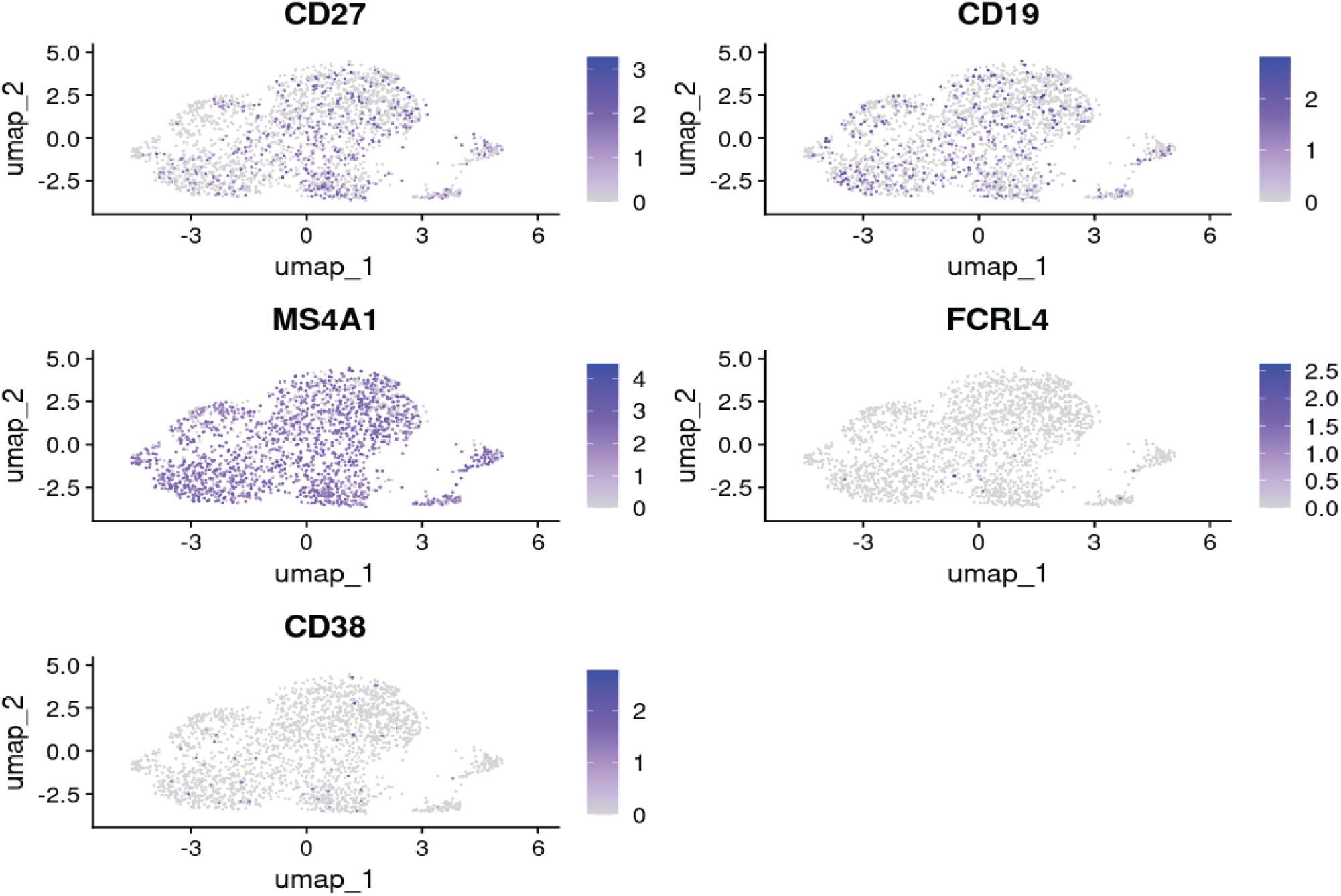
Expression of major genes of interest mapped to the UMAP representation of memory B cells from the normal lung tissue.

## Notes

### Competing Interest Statement

The authors have declared no competing interest.

### Summary of Updates

Section describing single cell analysis was revised in response to reviewers` questions. Figure 2 was revised for better image quality. Supplementary figures were added. Wording of the title was changed in response to reviewers comments.

